# Temporal *in vivo* platelet labelling in mice reveals age-dependent receptor expression and conservation of specific mRNAs

**DOI:** 10.1101/2021.12.17.473131

**Authors:** Paul C Armstrong, Harriet E Allan, Nicholas S Kirkby, Abhishek Joshi, Clemens Gutmann, Marilena Crescente, Jane A Mitchell, Manuel Mayr, Timothy D Warner

## Abstract

The proportion of young platelets, also known as newly formed or reticulated, within the overall platelet population has been clinically correlated with adverse cardiovascular outcomes. Our understanding of this is incomplete, however, because of limitations in the technical approaches available to study platelets of different ages. In this study we have developed and validated an *in vivo* ‘temporal labelling’ approach using injectable fluorescent anti-platelet antibodies to sub-divide platelets by age and assess differences in functional and molecular characteristics. With this approach we found that young platelets (<24h old) in comparison to older platelets respond to stimuli with greater calcium flux and degranulation, and contribute more to the formation of thrombi *in vitro* and *in vivo*. Sequential sampling confirmed this altered functionality to be independent of platelet size with no size differences or changes relative to the global population seen at any age. The age associated decrease in thrombotic function was accompanied by significant decreases in the surface expression of GPVI and CD31 (PECAM-1) and an increase in CD9. Platelet mRNA content also decreased with age but at different rates for individual mRNAs indicating apparent conservation of those encoding granule proteins. Our pulse-chase type approach to define circulating platelet age has allowed timely re-examination of commonly held beliefs regarding size and reactivity of young platelets whilst providing novel insights into the temporal regulation of receptor and protein expression. Overall, future application of this validated tool will inform on age-based platelet heterogeneity in physiology and disease.

## Introduction

Understanding the fundamental changes that occur during a platelet’s lifespan are central to many avenues of current research in both cardiovascular and hematological disease. However, research has been severely limited because of a lack of methodologies to identify circulating platelets of different ages. Platelets inherit a finite complement of messenger ribonucleic acid (mRNA) from the parent megakaryocyte^1^ which is rapidly lost once they are released into the circulation. As platelets are anucleate they are unable to replenish these levels^2^ and so nucleic acid dyes such as thiazole orange (TO) or SYTO13 have been widely used to differentiate young platelets based on their higher mRNA content (sometimes inferred to reflect an immature platelet fraction).^3–8^

Increased platelet turnover in a number of diseases, including diabetes mellitus, chronic kidney disease and essential thrombocythemia (ET)^9,10^ and the consequent increases in the ‘immature platelet fraction’ have been associated with a higher incidence of acute coronary syndromes^11^ and a reduced effectiveness of anti-platelet drugs.^12,13^ Similarly, several groups have utilized nucleic acid dyes and cell sorting to establish that young platelets defined in this way are hyper-reactive and of increased thrombotic potential.^4–6^ We have previously demonstrated that young platelets identified by higher mRNA content localize to the core of aggregates and drive aggregate formation even in presence of standard anti-platelet therapies, potentially reducing drug effectiveness in high platelet turnover conditions.^14^

Whilst useful as basic methodologies, there are limitations with mRNA dye-based approaches to fractionate platelets. Firstly, staining levels do not accurately advise on platelet age as fluorescence has not been temporally benchmarked. Secondly, following the rapid early loss of mRNA from young platelets the fluorescent signal can no longer discriminate platelets by ‘age’. Moreover, confounding factors including platelet size and nucleotide-rich granule content can cause undue bias or interference with counts or estimated fractions of immature young platelets. ^15,16^

As a result of these shortcomings there continues to be a search for alternative techniques to differentiate the newly formed platelet fraction. A common strategy in this regard has been to cause rapid thrombocytopenia in mice by injection of anti-platelet antibodies^17,18^. This acute depletion is followed by a burst of platelet production such that the entire circulating platelet population is taken as being reticulated or newly formed. However, such an approach involves a pathological insult to drive rapid platelet production and makes the questionable assumption that platelets released under such conditions are representative of platelets newly formed under normal circumstances.

We therefore sought to develop an alternative *in vivo* labelling approach in mice using antiplatelet fluorescent antibodies to provide a tool for the age-related tracking and isolation of platelet subpopulations. Here we demonstrate the application of this novel approach in revisiting widely held ideas regarding the phenotype of young platelets, identifying new age-related alterations in surface marker expression and revealing preferential retentions of mRNAs.

## Methods

### Ethical Statement

Animal procedures were conducted in accordance with Home Office approval (PPLs 70–7013 and PP1576048) under “The Animals (Scientific Procedures) Act” Amendment 2013 and were subject to local approval from Queen Mary University of London and Imperial College London Ethical Review Panels.

### Temporal labelling of platelets in vivo

*In vivo* grade fluorescently conjugated anti-platelet antibodies (anti-CD42c DyLight-x488 or x649, Emfret) were administered (*i.v*.) to the tail vein of C57BL/6 mice at specific timepoints as per supplier guidance. Labelling was confirmed and tracked using flow cytometry (Novocyte, Agilent Technologies) following micro-sampling from the tail vein. Comparative *in vitro* platelet labelling was achieved, where indicated using anti-CD41-BV421 (clone MWReg30, BioLegend).

### Blood collection and isolation of platelets

Mice were anesthetized with intraperitoneal (i.p.) ketamine (Narketan, 100 mg/kg; Vetoquinol) and xylazine (Rompun, 10 mg/kg; Bayer) and blood collected from inferior vena cava into lepirudin (Refludan, 25 μg/mL; Celgene, Windsor) or tri-sodium citrate (3.2%; Sigma). Platelet rich plasma (PRP) was isolated as previously published^19^. Briefly, whole blood was diluted 1:1 with modified Tyrode’s HEPES buffer (134mmol/L NaCl, 2.9mM KCl, 0.34mmol/L Na_2_HPO_4_, 12mmol/L NaHCO_3_, 20mmol/L HEPES and 1mmol/L MgCl_2_; pH 7.4; Sigma) and centrifuged at 100 *g*, 8 min.

### Immunofluorescence of temporally labelled platelets

Paraformaldehyde (4%; VWR) fixed platelets were centrifuged onto poly-L-lysine coverslips (VWR; 600 g, 5 min). Samples were mounted with Prolong Diamond antifade mount (Thermo Fisher Scientific). Confocal microscopy was performed using an inverted Zeiss LSM880 with Airyscan confocal microscope; 63× objective, 1.4 Oil DICII (Zeiss). Analysis was conducted using Zen Software (2.3 SP1, Zeiss) and ImageJ (NIH).

### Quantification of P-selectin expression and active confirmation of GPIIb/IIIa

PRP was diluted in phosphate buffered saline (PBS; 1:40) and incubated with protease-activated receptor-4 activating peptide AYPGKF amide (PAR4-amide, 200 μM; Bachem) or PBS for 20 min at 37°C in presence of anti-CD62P-BV421 (clone RB40.34; BD Bioscience) or anti-CD41/61-PE (clone JON/A, Emfret). Samples were subsequently fixed with 1% formalin and samples acquired and analyzed by flow cytometry (Novocyte) using NovoExpress software (Agilent Technologies).

### Quantification of calcium dynamics

Platelets were isolated from mice 24 hours after injection with anti-CD42c-DyLight-x649 and were subsequently incubated with Fluo-4 AM (5 μM; Thermo Fisher Scientific) for 30 min, 37°C, followed by counter staining with anti-CD41-BV421 (1:100; BD Biosciences) for 15 min and supplementation with calcium chloride (2 mM; Sigma). Baseline Fluo-4 fluorescence was recorded for 30 sec, followed by challenge with PAR4-amide (25 μM), collagen related peptide (1 μg/ml; CAMBCOL), the stable thromboxane (Tx) A_2_ mimetic U46619 (10 μM; Enzo) or ionomycin (10 μM; Invitrogen) and subsequent recording for 2 min. Samples were acquired on a BD LSRII (BD Bioscience) using FACSDiva acquisition software, and analyzed using FlowJo software v.10.

### *Ex vivo* whole blood aggregation

Platelet aggregation was performed in whole blood as previously described.^19^ Briefly, in half-area 96-well microtiter plates (Greiner Bio-One) whole blood was incubated with PBS, arachidonic acid (AA; 0.05–0.5mM; Sigma), Horm collagen (0.1–10μg/ml; Takeda), PAR4-amide, (50–100μM), or U46619 (0.1–10μM) under mixing conditions (1200rpm, Bioshake iQ, Qinstruments). Samples were subsequently diluted (1:20 acid citrate dextrose) and analyzed by flow cytometry (Novocyte) to identify and determine proportions of the single platelet population.

### Pulmonary embolism injury model

Pulmonary embolism (PE) was induced as previously described.^20^ Briefly, Horm collagen (50μg/kg, Takeda) and U46619 (210μg/kg, Enzo) were administered i.v. Blood was microsampled before and after (4 min) administration of platelet activators for determination of circulating labelled platelets.

### Surface marker expression

PRP obtained from temporally labelled mice was diluted 1:40 in modified Tyrode’s HEPES buffer and incubated 20 min at room temperature with one of the following PE-conjugated antibodies: anti-CD9 (clone MZ3), anti-CD49b (clone DX5), anti-CD31 (clone 390), anti-H2 (clone 1/42; all BioLegend) or anti-GPVI (clone 784808, R&D systems). PE-conjugated isotype antibodies of Rat IgM (clone RTK2118), IgG1 (clone 2071) or IgG2a (clone RTK2758; all BioLegend) were used as corresponding non-specific binding controls. After staining, samples were fixed with 1% formalin and acquired on Novocyte flow cytometry and analyzed using FlowJo v10 software.

### RNA extraction, cDNA synthesis and quantitative real time polymerase chain reaction (qRT-PCR) of sorted, temporally labelled platelets

Full details of mRNA analysis of platelet populations are available in the supplementary methods. RNA was extracted from FACS-sorted platelets (2.5 million per population), using the miRNAEasy mini kit (Qiagen, 217004), following manufacturer’s instructions. Samples were spiked using Cel-miR-39-3p (Qiagen, 219600) and MS2 carrier RNA (Roche, 10165948001) as controls and cDNA was generated using VILO RT Superscript (Thermo Fisher, 11755-250) from 8μL of RNA, as per manufacturer’s instructions. Transcripts were quantified using TaqMan qPCR RNA primers for selected targets, in a ViiA 7 Real-Time PCR System (Applied Biosystems) using ViiA 7 Software (Applied Biosystems).

### Statistics and data analysis

Data is presented as mean ± standard error of the mean (SEM), and analyzed using FlowJo v10 (Treestar) and Prism v9 (GraphPad software, USA). For analysis, the “single platelet” population was gated based on forward scatter and anti-platelet immunoreactivity (fluorescence intensity). Statistical significance was determined by two-way ANOVA with Dunnett’s post-hoc test unless otherwise stated, and data sets considered different if p<0.05. Each n value represents a data point from a separate animal.

## Results

### Temporal fluorescence labelling of platelets in vivo to identify age-defined populations

Intravenous injections of short-lived fluorescence-labelled anti-CD42c antibody (DyLight-x488) produced ~95% labelling of circulating platelets (supplemental Figure S1) that remained fluorescently tagged for the duration of their life. After 24 hours, an un-labelled platelet subpopulation, representing approximately 20% of the entire population, was seen as a result of normal platelet removal and replacement. At this time an injection of the same anti-CD42c antibody conjugated to an alternative fluorophore (DyLight-x649) created two differentially fluorescently-labelled platelet sub-populations (Figure 1A). A minority single labelled (x649+/x488-) population corresponding to platelets <24h old and a majority dual-labeled (x649+/x488+; Figure 1B) population corresponding to platelets >24h old.

**Figure 1:**
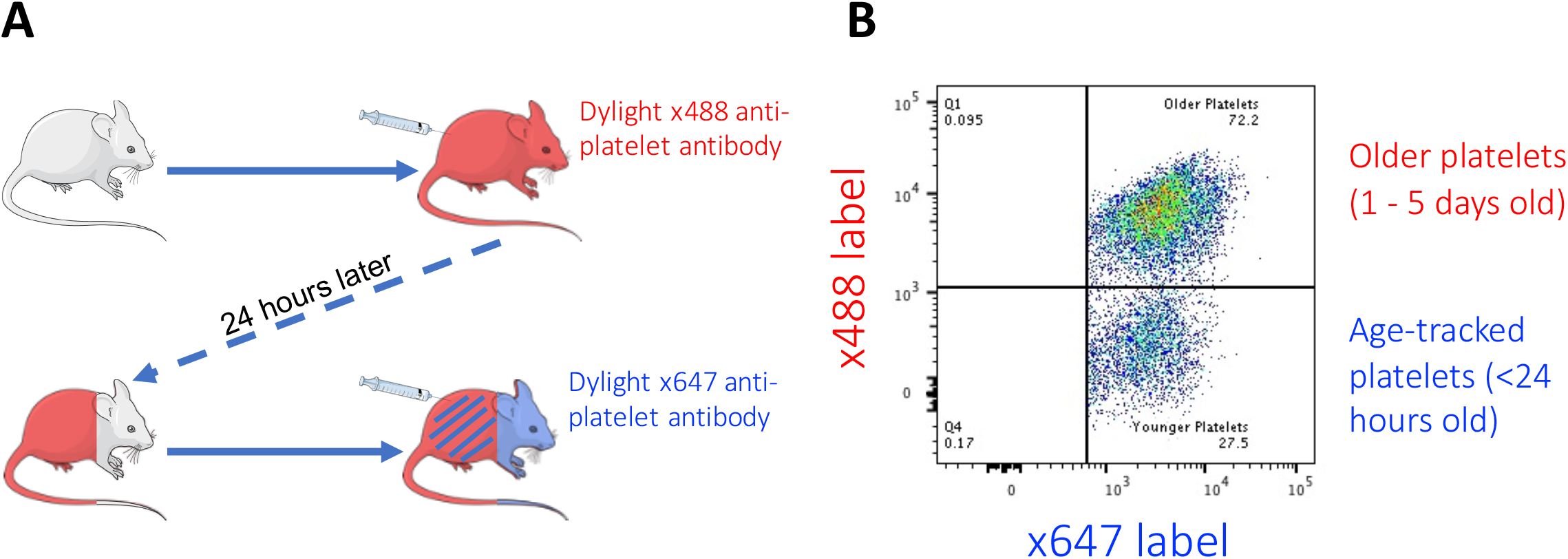
Methodological diagram of *in vivo* temporal labelling of platelets in mice using fluorescently-tagged antibodies. Mice are injected with Dylight x488 labelled anti-CD42c to achieve near universal labelling (>95%) followed by a second injection after 24 hours with Dylight x649 labelled anti-CD42c results in dual labelled platelet population and a minority x649 single label population corresponding to platelets generated in intervening 24-hour period since first injection. Representative pseudo-colour dot-plot of differentially-labelled platelets from blood sample as detected by flow cytometry.

### Younger platelets demonstrate increased reactivity, contributing disproportionately to thrombotic response

No significant differences were observed in basal P-selectin expression or JON-A binding between platelets <24h and >24h old (Figure 2A, B). Following incubation with a PAR-4 peptide agonist, the younger platelets demonstrated higher P-selectin expression (83147±858 MFI) than older platelets (80880±1257 MFI; p<0.005; Figure 2A) but lower JON-A binding (864±18 vs 884±17; p<0.01; Figure 2B). Younger platelets also displayed significantly stronger increases in intracellular calcium levels following exposure to PAR4 peptide (Figure 2C), ionomycin (Figure 2D) and CRP-XL or U46619 (supplemental Figure S2).

**Figure 2:**
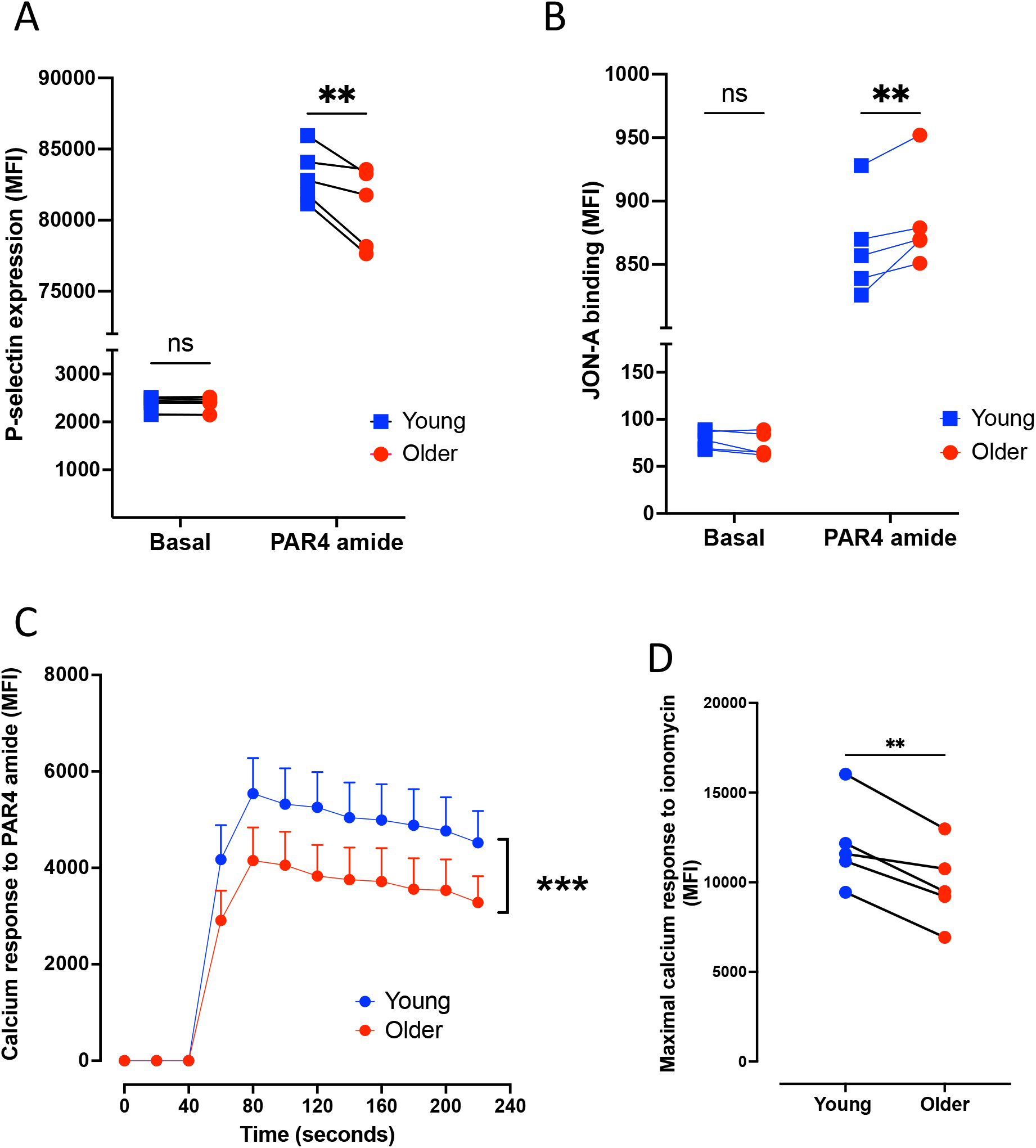
Differential degranulation, integrin activation and calcium flux of young platelets compared to older platelets. Younger (<24h old) platelets express (A) higher levels of P-selectin but (B) lower levels of active GPIIb/IIIa in response to PAR4-specific stimulation compared to corresponding older platelets in the same sample. (C) Younger platelets have a greater calcium flux in response to PAR4 specific stimulation (n=5) consistent with a (D) greater maximal calcium response in response to ionomycin. Data are individual replicates or mean ± SEM (C). **p<0.01, ***p<0.001 by two-way ANOVA or paired t-test as appropriate.

Next, to assess the relative aggregatory potential of young and older platelets, the proportions of each sub-population were determined in whole blood samples following stimulation. In such experiments equal aggregatory potential and contribution would result in no differences between the proportions found in the un-aggregated single platelet population before and after stimulation. However, in response to multiple agonists, the proportions diverged significantly. In response to PAR4 peptide the proportion of younger platelets remaining unaggregated significantly decreased (24.7±0.3% to 18.6±0.6%, p<0.05) whilst the older population significantly increased (75.3±0.3% to 81.4±0.6%, p<0.05; Figure 3A). Similarly patterns of response were seen when aggregation was stimulated by CRP-XL or U46619 (supplemental Figure S3). To confirm these *in vitro* observations *in vivo*, we employed a murine model of pulmonary embolism and assessed circulating proportions of the labelled platelet populations pre- and post-embolism. Compared to pre-injury, a significant decrease in the circulating proportion of young platelets (19.0±1.7% to 17.0±1.8%, p<0.05) and a corresponding significant increase in the proportion of older platelets remaining unaggregated (80.2 ±1.8% to 82.9 ±1.7%, p<0.05, Figure 3B) was observed.

**Figure 3:**
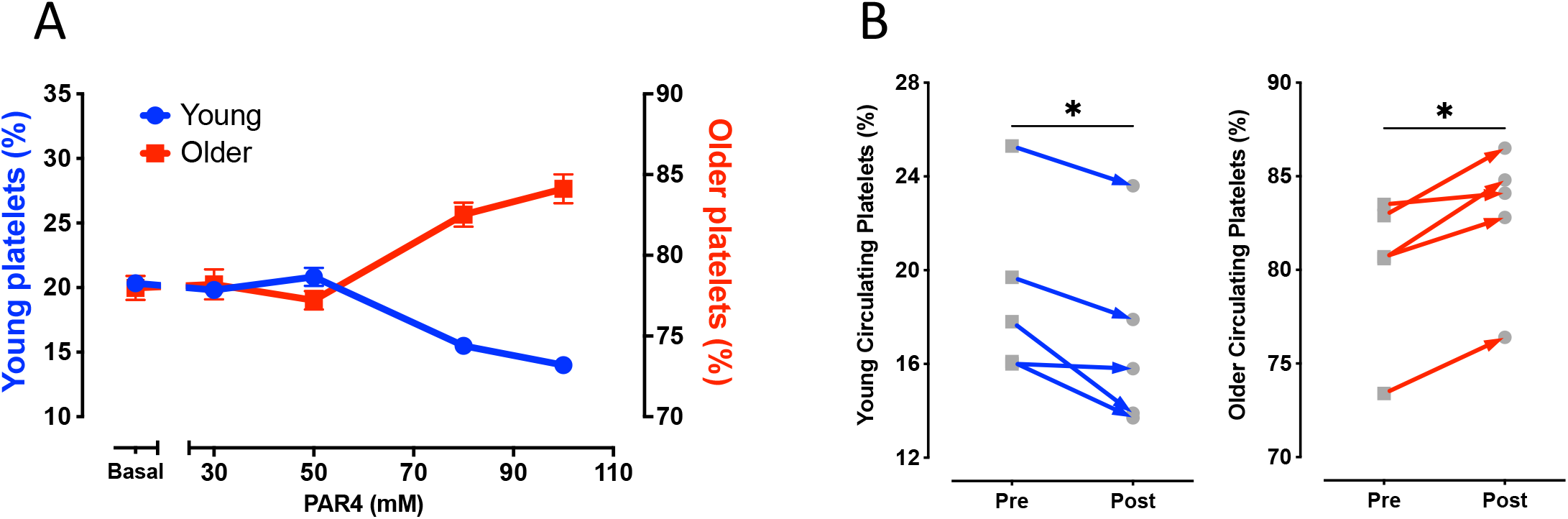
Younger platelets disproportionately contribute to thrombotic response. (A) Population tracking in whole blood following stimulation with increasing concentrations of PAR4-specific stimulation to drive *ex vivo* aggregation resulted in decreasing proportions of younger platelets and corresponding increased proportions of older platelets remaining in the non-aggregated single platelet portion. (B) Circulating proportions of labelled younger platelets *in vivo* decreased following pulmonary embolism challenge with corresponding increase in proportions of circulating older platelets. Data are mean ± SEM (A) or individual replicates. *p<0.05, **p<0.01 by two-way ANOVA or paired t-test as appropriate.

### Newly formed platelets are heterogenous in size and do not change size distribution with age

Platelets identified as <24h old were heterogenous in size with no discernable difference in size distribution to the global platelet population as determined using cytometry derived forward- and side-scatter (FSC and SSC respectively). The same tracked-population was heterogenous and similarly sized relative to the global population in samples obtained from the same mice across days 2-5. Notably the size distribution of the tracked subpopulation did not change as it aged in the circulation (Figure 4A, supplemental Figure S4). Furthermore, immunofluorescence confirmed there was no difference in platelet cross sectional area between young (<24h) and older (1-5 days) platelets (5.28±0.13μm vs. 5.24±0.25μm respectively; p>0.05; Figure 4B, C).

**Figure 4:**
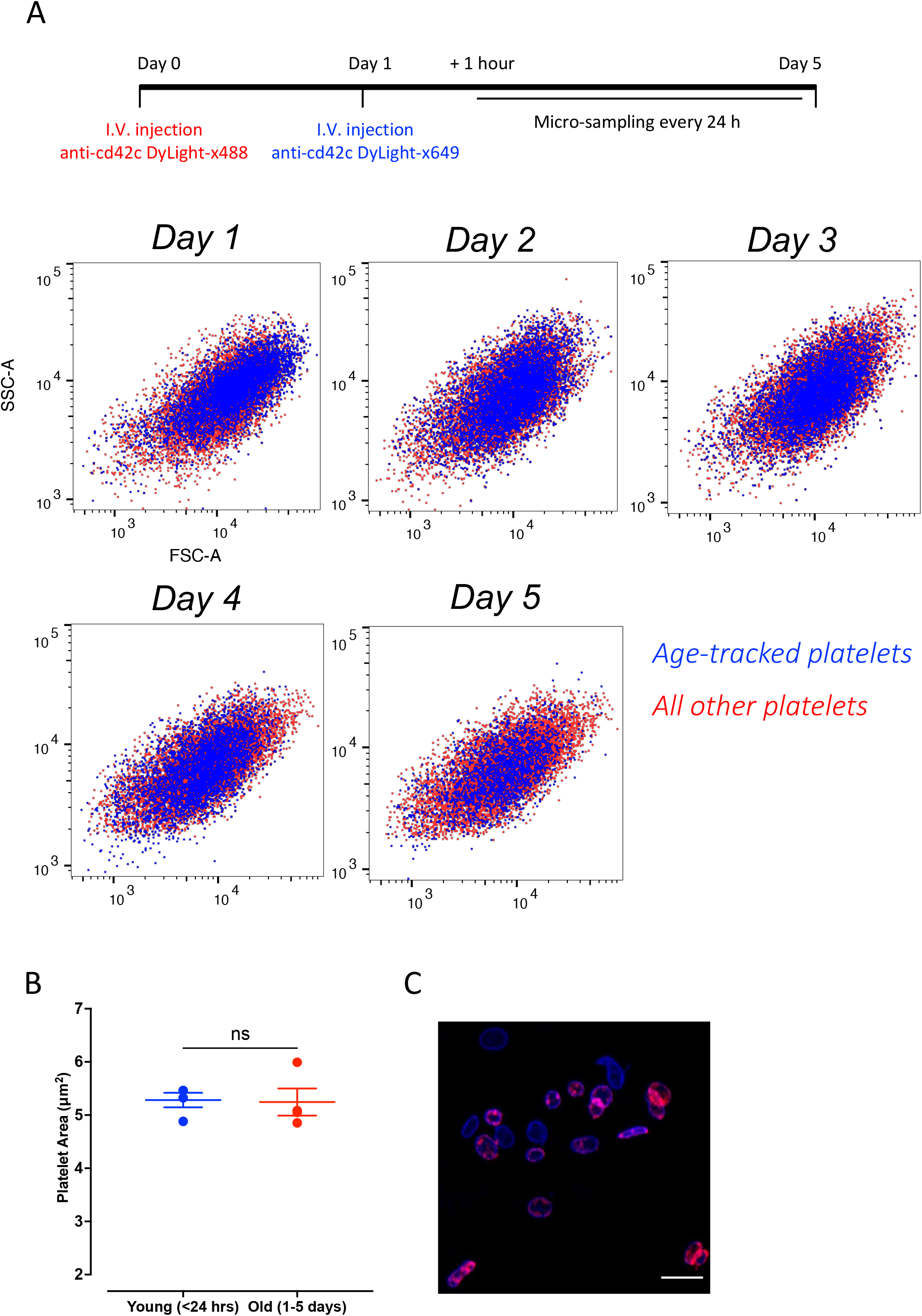
Platelets are heterogenous in size when formed and do not change size with age in the circulation. Following temporal-labelling, serial blood samples were obtained each day for 5 days. (A) Representative data of ‘Age-tracked’ platelets (blue) in an individual mouse. Tracked platelets were detectable in circulation for the full 5-days and were comparable in size to the counter-stained entire platelet population (red) as determined by flow cytometric forward scatter (FSC) and side scatter (SSC) properties. No change in size for the tracked population was observed throughout sampling time course. (C) Platelet area measured by confocal microscopy confirmed equal size of younger and older platelets. (D) Representative image of younger (blue) and older (blue & red) present in analyzed platelet samples isolated from labelled mouse. Data are individual replicates with overlaid mean ± SEM and analyzed by paired t-test.

### Major Histocompatibility class (MHC) I expression decreases with platelet age

MHC-I expression on platelets aged <24h was higher than the global median and decreased significantly with age (Figure 5A) equating to a loss of 30±6% across the platelet lifespan.

**Figure 5:**
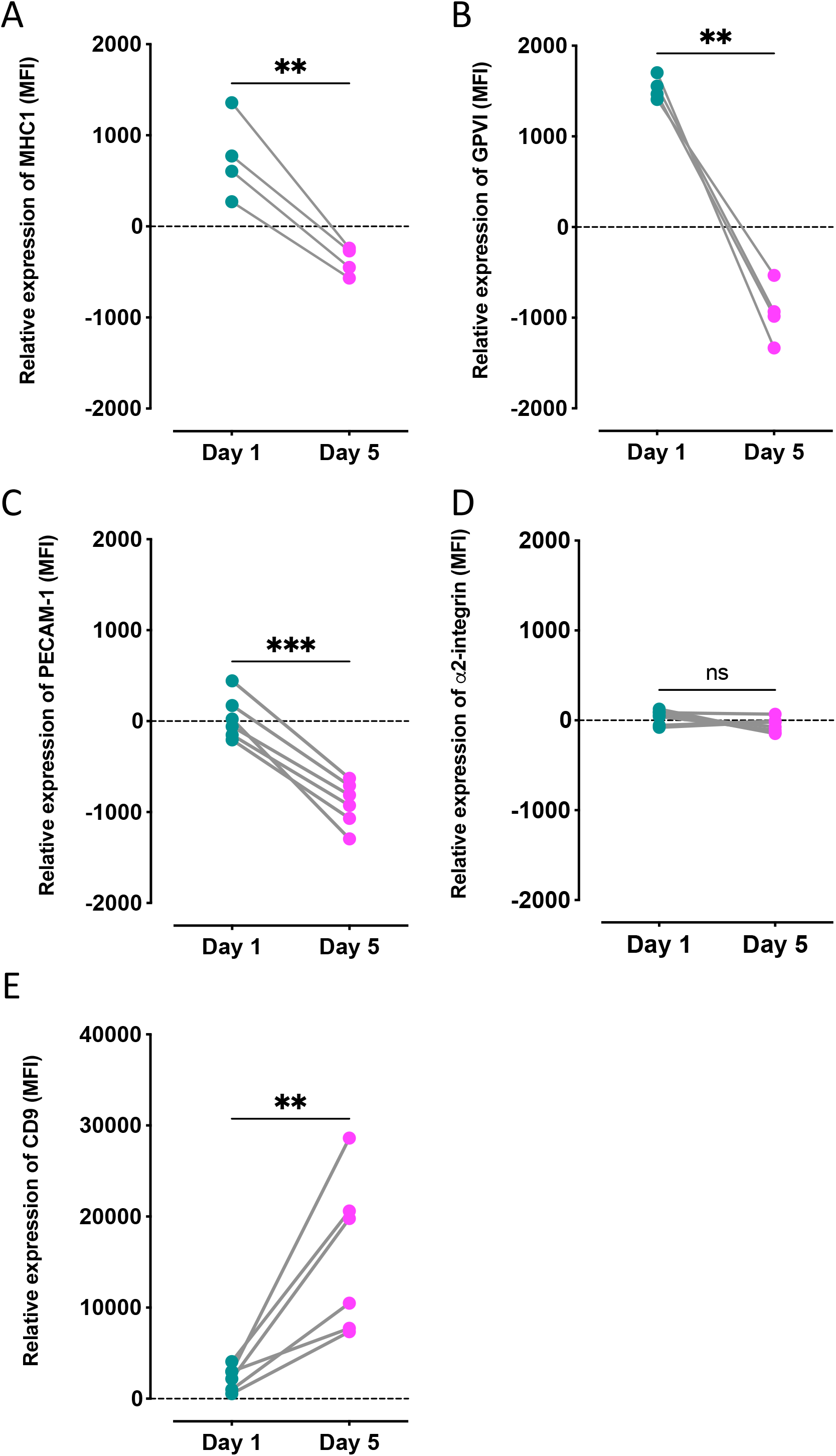
Differential surface receptor expressions of ageing platelets in circulation. Expression of receptors on age-tracked platelets, relative to global population expression, determined by flow cytometry in samples obtained from individual mice on day 1 (green) and 5 (cyan) post labelling. Significant decreases in expression of (A) MHC-1, (B) GPVI and (C) PECAM-1 between day 1 and day 5 of age. (D) No difference in expression of α2 integrin but (E) significant increase in CD9 expression. **p<0.01, ***p<0.001 as analyzed by paired t-test.

### Platelets lose GPVI and PECAM-1 but increase expression of CD9 with age

GPVI expression was significantly higher on <24h old platelets, and significantly lower at day 5 when compared to the global medians, a total loss equating to 39±2% (p<0.01, Figure 5B). Expression of CD31 significantly decreased between days 1 and 5 (20±3% loss, p<0.001; Figure 5C). In contrast, CD9 expression was significantly higher than the global median on day 5 (p<0.01, Figure 5E) equating to an increase of 13±4% relative to day 1. No significant changes in expression of CD49b were found (Figure 5D).

### Platelets favor retention of granule protein encoding mRNA

Principal component analysis of mRNA content demonstrated three analyzed populations, younger (<24h), older (1 -5 days old) and global, clustered individually (Figure 6A). The loading plot of individual mRNAs demonstrated a unidirectional pattern for PC1 indicating that a decreasing abundance of mRNAs likely predominates this algorithm. Interestingly, however, mRNAs also appeared to cluster separately according to PC2 indicating potential for individual patterns of decreased abundance (Figure 6B). Further investigation grouping mRNAs by function, namely signaling, structural, receptors and granules, suggested again a reduction in abundance associated with age, with <24h old platelets having the highest and 1–5-day old platelets having the lowest amounts of each mRNA (Figure 6C). However, the amount of loss varied by specific mRNA (median fold change ranging from −5.3 to −66.4). These changes were not contingent on starting levels (Figure 6D) but there was some association with functional categorization. In particular, those mRNAs classified as associated with signaling proteins demonstrated the greatest reduction in abundance whilst those associated with granule proteins demonstrated the least (median fold change: −36.8 versus −8.9 respectively).

**Figure 6:**
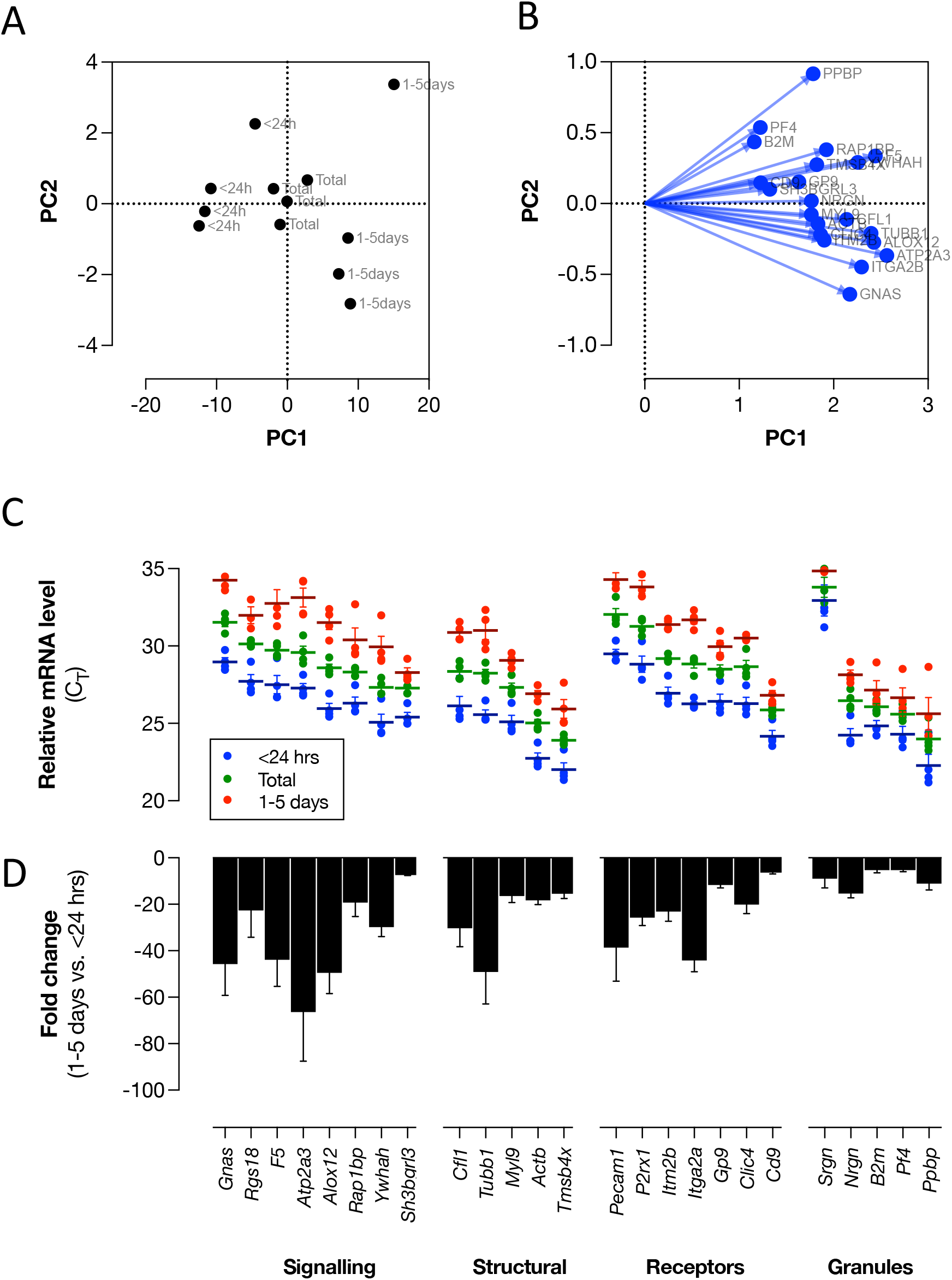
Platelet mRNA content decreases with age but differential loss according to function of encoded protein. Younger, older and total platelet populations were isolated by FACS and mRNA extracted for comparative analysis. (A) PCA analysis of samples resulted in individual clustering of samples according to age with (B) loadings analysis depict unidirectional relationship for PC1 but differential PC2 relationship. Grouping of measured mRNAs by downstream function and (C) relative mRNA level analysis illustrates variable content of individual mRNAs but with universal loss from highest levels in younger (blue) to total (green) to lowest levels in older (red) platelets. (D) Calculated fold-change from younger to older populations reveals differential scales of loss within groupings with mRNAs associated with encoding granule proteins experiencing greatest retention. Data are mean ± SEM (n=4).

## Discussion

We have developed an *in vivo* labelling approach which allows for age-related sub-populations of platelets to be identified and tracked. This technique can be coupled with a range of functional and molecular analyses to provide unique insights into platelet age-associated physiology. Here we have used it to address the two fundamental assumptions that newly formed platelets are more reactive and bigger. Furthermore, we have used this new approach to gain insights into changes in the expression of platelet receptors and the retention of mRNA during ageing that may underpin alterations in their functionality.

The nature of the relationship between platelet age, size and reactivity has remained contentious for many years. Perhaps the point of least contention is that mRNA-high ‘young’ platelets are more reactive, with a raft of literature over the years using *in vitro, ex vivo* and *in vivo* studies supporting this hypothesis^14,21^. Indeed, our recent studies identifying platelet age through RNA labeling of healthy human platelets are consistent with a decline in hemostatic function as platelets age. We demonstrated that young platelets express higher levels of P-selectin and have an enhanced calcium signaling capacity.^4,6^ Supporting our work in healthy human blood, our temporal labelling approach confirmed the youngest platelets have enhanced degranulation and calcium signaling in response to a range of agonists. Conversely, following stimulation older platelets have increased levels of the activated GPIIb/IIIa receptor, indicating a higher propensity for fibrinogen binding. Despite this, *ex vivo* aggregation studies and an *in vivo* PEi thrombosis model demonstrate that under healthy turnover conditions young platelets contribute highly and disproportionately to forming thrombi.

A factor that continues to confound our understanding of why young platelets are more reactive, or have higher thrombotic potential, is the enduring dogma that young platelets are larger. Originally proposed in the 1960s^22^ this hypothesis argues that younger platelets have greater volume resulting in greater granule content, and greater surface area resulting in greater surface receptor number. In support of this hypothesis, *in vivo* platelet studies in a range of species following pathological insult such as platelet-depletion have observed that the platelets acutely formed in response are larger in size.^23^ Similarly, associations have been found between mean platelet volume and immature platelet fraction measurements in patient populations. However, a series of papers in the mid 1980s reported no correlation between platelet age and size.^24–26^ Similarly, *in vivo* biotinylation studies in healthy rabbits have shown that younger platelets are no larger than older platelets.^27^ We therefore coupled our temporal labelling technique to flow cytometry and immunofluorescence to assess the sizes of platelets of different ages. Our data clearly shows that in healthy mice younger and older platelets do not differ in mean size or size distribution. Notably, the aforementioned *in vivo* studies that associate younger platelets with larger size are predicated on a pathological insult and the subsequent acute response to restore platelet number. It seems reasonable to conclude that the processes that underpin such an acute response differ from those supporting normal platelet production and release and could produce a different population of platelets.

Similar research has also been applied to identifying other markers of age. If platelet size does not change with age, coupled with our recent observation of loss of protein content in ageing human platelets^6^ might suggest support proposals that platelet density is a good marker of platelet age^28^. However, the calculation of platelet density is technically challenging and laborious whereas the measurement of surface molecular markers or receptor expression is much more easily accomplished. Angenieux *et al*^29^ recently identified that across the platelet population, young platelets, identified by thiazole orange staining, had the highest levels of MHC-1/HLA-1. In our study using defined temporal labelling we confirmed this observation and found novel differential expression patterns according to platelet age for several other receptors associated with collagen responsiveness. We identified higher surface expression levels of GPVI and CD31 in younger compared to older platelets, whilst conversely the levels of CD9 increased as platelets age in the circulation. The origin of this increase in surface CD9 is unclear but de novo production or accumulation from exosomes^30^ are two possibilities. Despite its functional role remaining unclear^31^, CD9 expression is generally high on platelets^32^ and it has been shown to localize with GPVI^33^ and ADAM10^34^ within tetraspanin-enriched microdomains. Both GPVI and CD31 undergo proteolytic cleavage mediated by ADAM-10 and MMP2 respectively under physiological and pathological conditions^35–37^, and therefore constitutive or triggered activity may contribute to their decline in surface expression as platelets age. With relevance to the decline in platelet responsiveness associated with ageing, a decrease in GPVI expression would correlate with the impaired CRP-XL-induced activation observed in our functional studies.

In addition to changes in receptor expression, we sought to examine the mRNA content of platelets. Young platelets are well accepted as having the highest levels of mRNA and that this mRNA content decreases with time.^38^ Indeed the results of our measurement, from the selection of mRNA, would support this view. Platelets have been reported as containing ~10,000 different mRNA transcripts^39^ and separately that this mRNA has a half-life of ~6 hours^38,40^. We wished to examine if this loss of mRNA was universal using our temporal labelling approach. Unbiased principal component analysis confirmed that young platelets had higher overall levels of mRNA but also revealed a differential reduction in specific platelet mRNAs. Further analysis revealed the rate of loss for mRNAs to vary greatly with those encoding granule proteins experiencing the least reduction. This variation in rates of loss strongly suggests that previous reports of 6 hours as the decay half-life of platelet mRNA^38,40^ is a general number while the decay of particular mRNAs is under more refined control. The molecular nature and purpose of this apparent conservation will be important to elucidate but research by Mills *et al*^40^ has indicated that ribosomes protect mRNA through binding which may enhance specific translation. Following our observation that mRNAs that encode granule proteins are particularly conserved it is possible to hypothesize that this will support their specific protein synthesis. *De novo* protein synthesis in platelet is not a new concept as it has been reported for over 50 years^41^ and in particular in response to activation^42–44^. However due to the technically challenging nature of these studies much interrogation of this process has occurred in an *ex vivo* setting and questions remain over the quantity of proteins produced^45^ and their physiological and/or pathophysiological significance. Our labelling approach will help to unravel whether such de novo synthesis is part of a maturation step of platelets *in vivo* or, following reports of selective protein packaging and release of granular contents and extracellular vesicles^46–48^, whether selective retention of granular mRNA for protein synthesis prolongs their period of reactivity.

In summary, we present a novel approach for discovering age-related alterations in platelet form and function. Our approach has discovered changes in platelet GPVI, CD31 and CD9 expression as platelets age in the circulation, and also provides insights into differential regulation and retention of platelet mRNAs. These changes may well underpin alterations in platelet functionality during the ageing process. Therefore, this study reports novel insights into fundamental assumptions of platelet size and reactivity as they age in the circulation and provides an approach that will act as an important tool for future research.

## Acknowledgements

This work was supported by the British Heart Foundation (RG/19/8/34500 to TDW and PCA, FS/18/60/34181 to CG and FS/16/31699 to NSK) and the Wellcome Trust (101604/Z/13/Z).

## Authorship Contributions

PCA designed the research, performed the assays and collected data, analyzed and interpreted data, performed statistical analysis, and wrote the manuscript. HEA performed assays, analyzed, and interpreted data, performed statistical analysis, and revised the manuscript. AJ and CG performed analysis, and collected and analyzed data, MC assisted with the assays and collecting data, NSK, JAM, MM and TDW analyzed and interpreted data, and revised the manuscript.

