## Supplemental Figures for "Temporal *in vivo* platelet labelling in mice reveals age-dependent receptor expression and conservation of specific mRNAs"

**Short title:** Temporal mouse platelet labelling

**Corresponding Author:** Paul Armstrong, Centre for Immunobiology, Blizard Institute, 4 Newark Street, Barts and the London School of Medicine and Dentistry, Queen Mary University of London, London, E1 2AT, UK

### Supplemental Figures

### Supplemental Figure S1

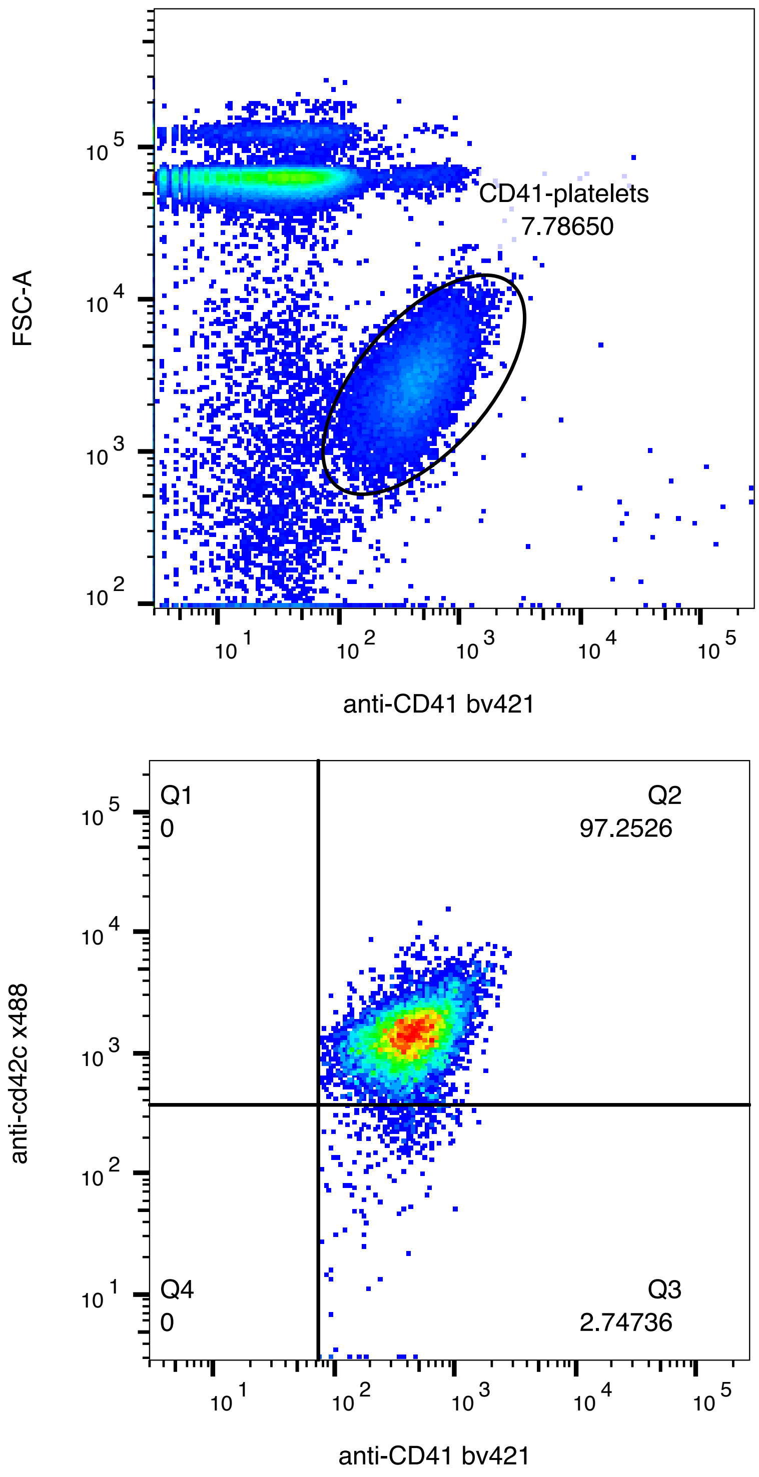

### Supplemental Figure S2

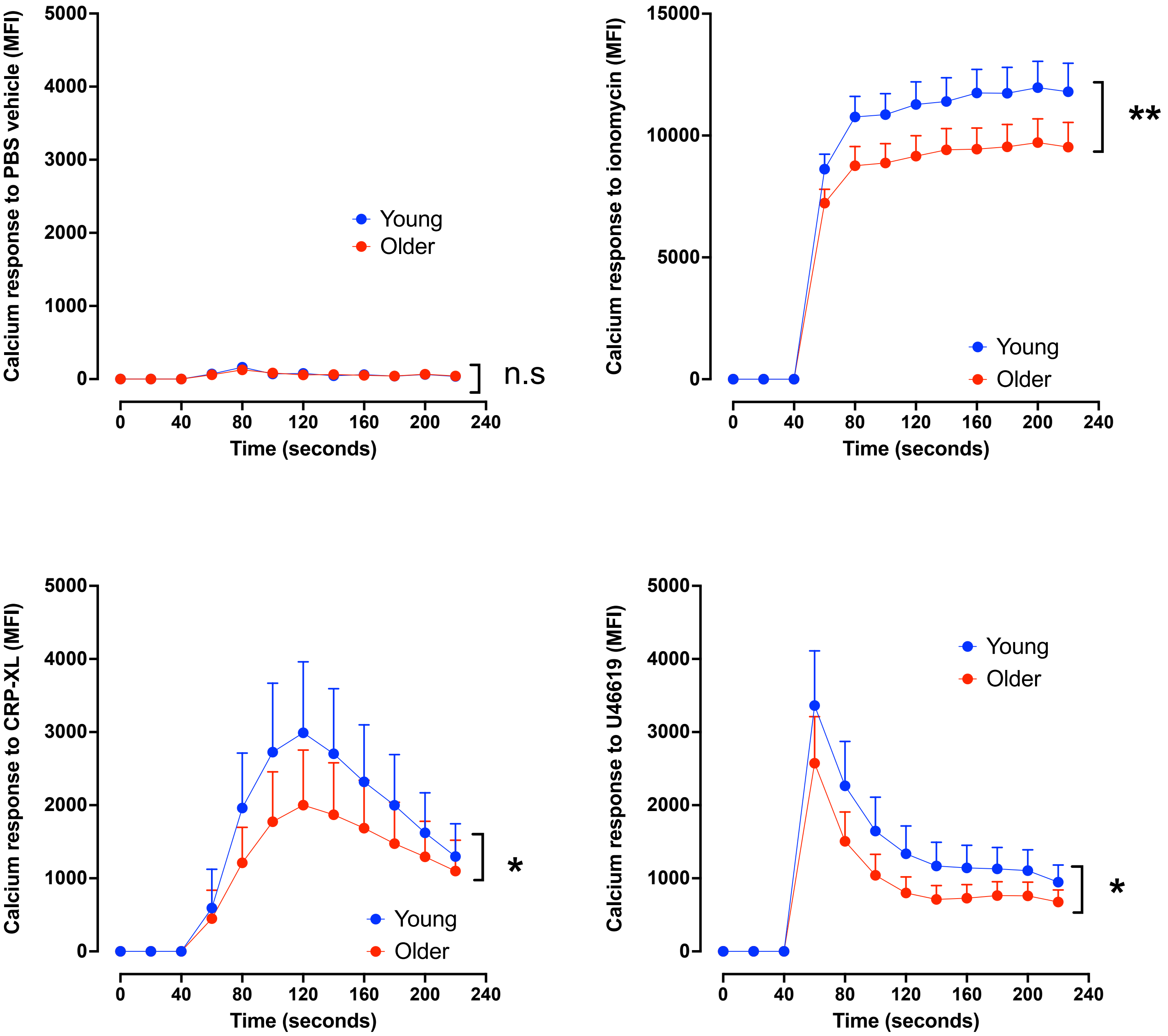

Supplemental Figure S3

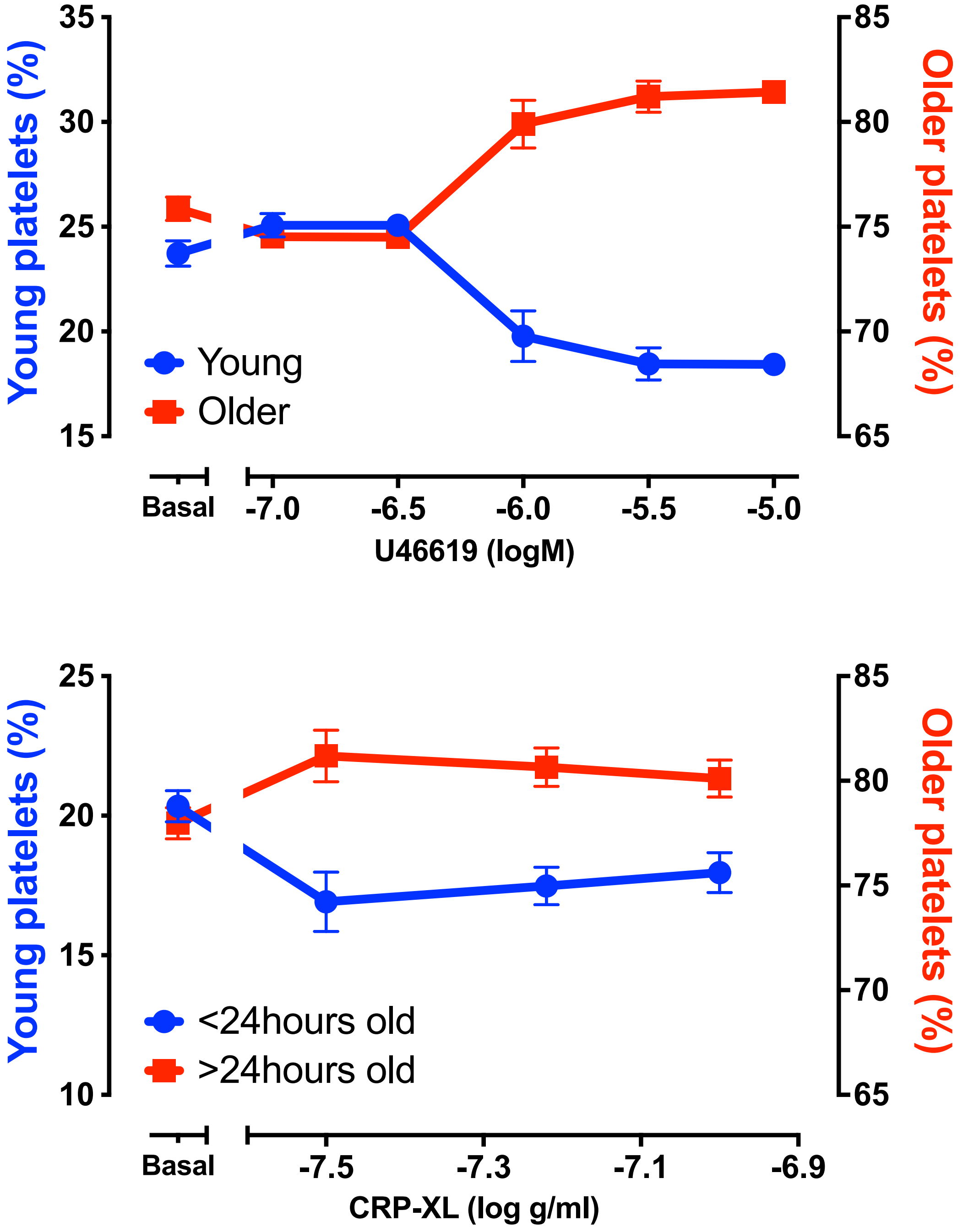

### Supplemental Figure S4

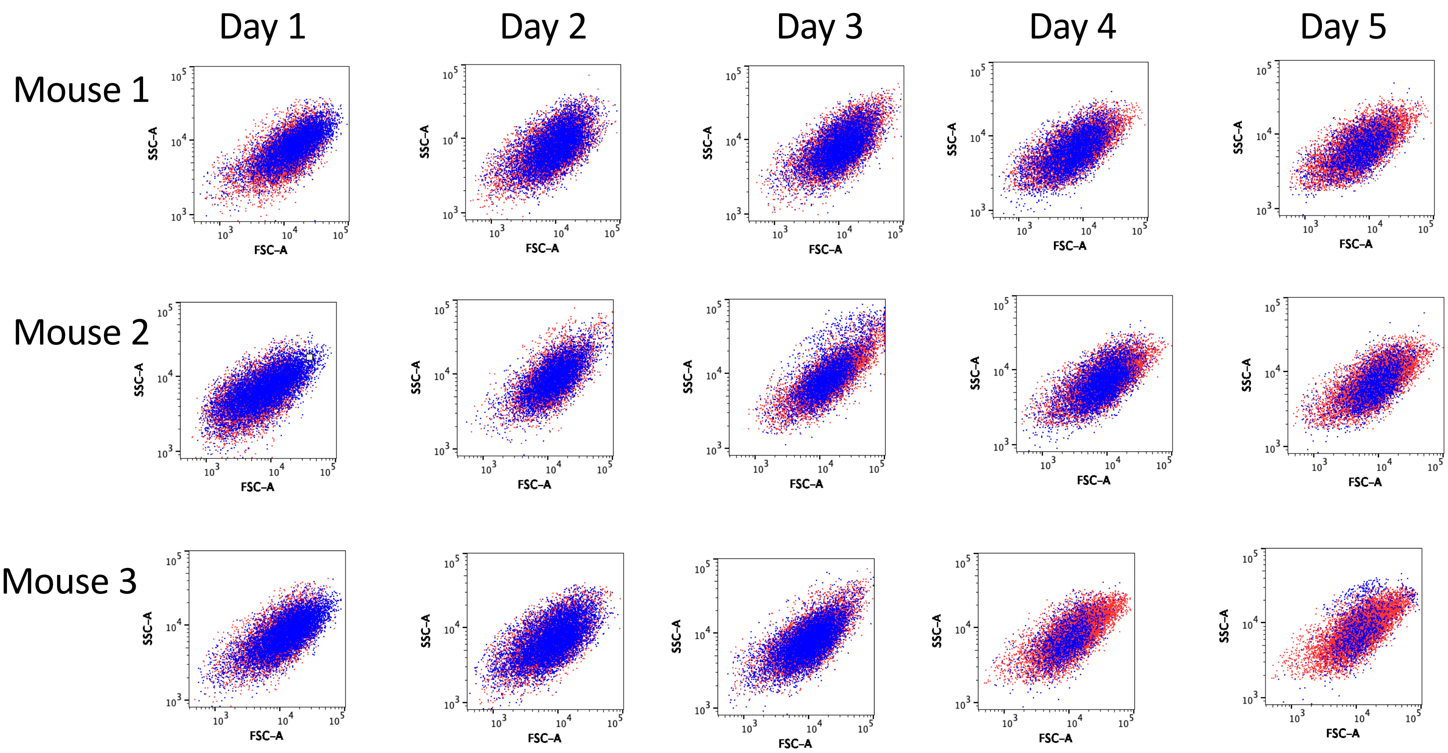
